# Identification of Stem Cells from Large Cell Populations with Topological Scoring

**DOI:** 10.1101/2020.04.08.032102

**Authors:** Mihaela E. Sardiu, Box C. Andrew, Jeff Haug, Michael P. Washburn

**Author notes:** To whom correspondence should be addressed Contact: Michael P. Washburn, Ph.D., Stowers Institute for Medical Research,1000 E. 50th St, Kansas City, MO 64110. Authors contributed equally to this work (Mihaela E. Sardiu, Box C. Andrew).

## Abstract

Machine learning and topological analysis methods are becoming increasingly used on various large-scale omics datasets. Modern high dimensional flow cytometry data sets share many features with other omics datasets like genomics and proteomics. For example, genomics or proteomics datasets can be sparse and have high dimensionality, and flow cytometry datasets can also share these features. This makes flow cytometry data potentially a suitable candidate for employing machine learning and topological scoring strategies, for example, to gain novel insights into patterns within the data. We have previously developed the Topological Score (TopS) and implemented it for the analysis of quantitative protein interaction network datasets. Here we show that the TopS approach for large scale data analysis is applicable to the analysis of a previously described flow cytometry sorted human hematopoietic stem cell dataset. We demonstrate that TopS is capable of effectively sorting this dataset into cell populations and identify rare cell populations. We demonstrate the utility of TopS when coupled with multiple approaches including topological data analysis, X-shift clustering, and t-Distributed Stochastic Neighbor Embedding (t-SNE). Our results suggest that TopS could be effectively used to analyze large scale flow cytometry datasets to find rare cell populations.

## Introduction

Utilizing high-throughput technologies, dynamic -omics data including genomics, transcriptomics, epigenomics, proteomics, and metabolomics has produced temporal-spatial big biological datasets which generally can be analyzed using similar approaches ^1, 2^. Statistically, - omics data is typically presented as a large data matrix where the rows correspond to variables like the expression level of a gene in genomics, the expression level of a protein in proteomics, and the expression of protein markers on a cell in flow cytometry, and the columns correspond to independent samples ^3, 4^. Major challenges persist regarding the analysis of large scale -omic datasets. This includes challenges regarding how to handle the complexity of data and how should the data be translated to discover the underlying biology from these large and complex matrices.

It is therefore necessary to use different analysis methods or scoring strategies for large scale datasets to achieve more biological understanding and generate novel hypotheses. We recently introduced a new topological score for the analysis of proteomics data named Topological Scoring (TopS) ^5, 6^. The TopS method has already been used in an analysis of biological networks and its performance has been tested against other tools for proteomics analysis ^5-8^. TopS uses a likelihood score on quantitative values and in principle is can use any type of quantitative data, rather than being restricted to one type of -omics data. TopS generates large and small values corresponding to strong or weak links between variables and samples relative to other samples in a matrix ^5, 6^. In general, TopS in combination with machine learning can be used to detect subnetworks consisting of points with similar patterns in large networks.

Flow cytometry is a technology that typically generates large scale quantitative datasets for the discovery of specific or rare cell populations such as bone marrow-residing hematopoietic stem cells (HSCs) ^9, 10^. The ability to detect specific cell populations that associate strongly with different *cell-surface* protein markers typically presents challenge to data analysis and many clustering methods have been used to study such a dataset) ^9, 10^. Here, we report the results of analyzing the *Nilsson rare* human hematopoietic stem cell dataset ^9, 10^ set by TopS and machine learning (Figure 1). We compared the use of TopS to original transformed data and expert gating results to test the usage of TopS for the analysis of a multi-color cytometry data set. Here we implement three different computational approaches-based on machine learning including topological data analysis (TDA) ^11-14^, X-shift clustering ^15-17^, and t-Distributed Stochastic Neighbor Embedding (t-SNE) ^18-20^ analysis for the analysis and visualization of the flow cytometry data. TDA is one of the newer and powerful method for the analysis of large datasets ^11-14^. TDA is using topological and geometric approaches to infer relevant features in complex datasets. X-shift clustering has been used in the analyses of the CyTOF and flow cytometry datasets and it is using a weighted K-nearest neighbor density estimation (KNN-DE) to determine the clusters in a large dataset ^15-17^. Lastly, t-SNE is a non-linear technique for dimensionality reduction that is commonly used for the visualization of high-dimensional datasets ^18-20^. Unlike TDA and X-shift, t-SNE is often used with other unsupervised learning algorithms for data classification. We demonstrate that TopS is an effective approach for processing data prior to utilization of TDA, X-shit, or t-SNE and is capable of efficiently finding rare cell populations in a flow cytometry sorted human hematopoietic stem cell dataset

**Figure 1.**
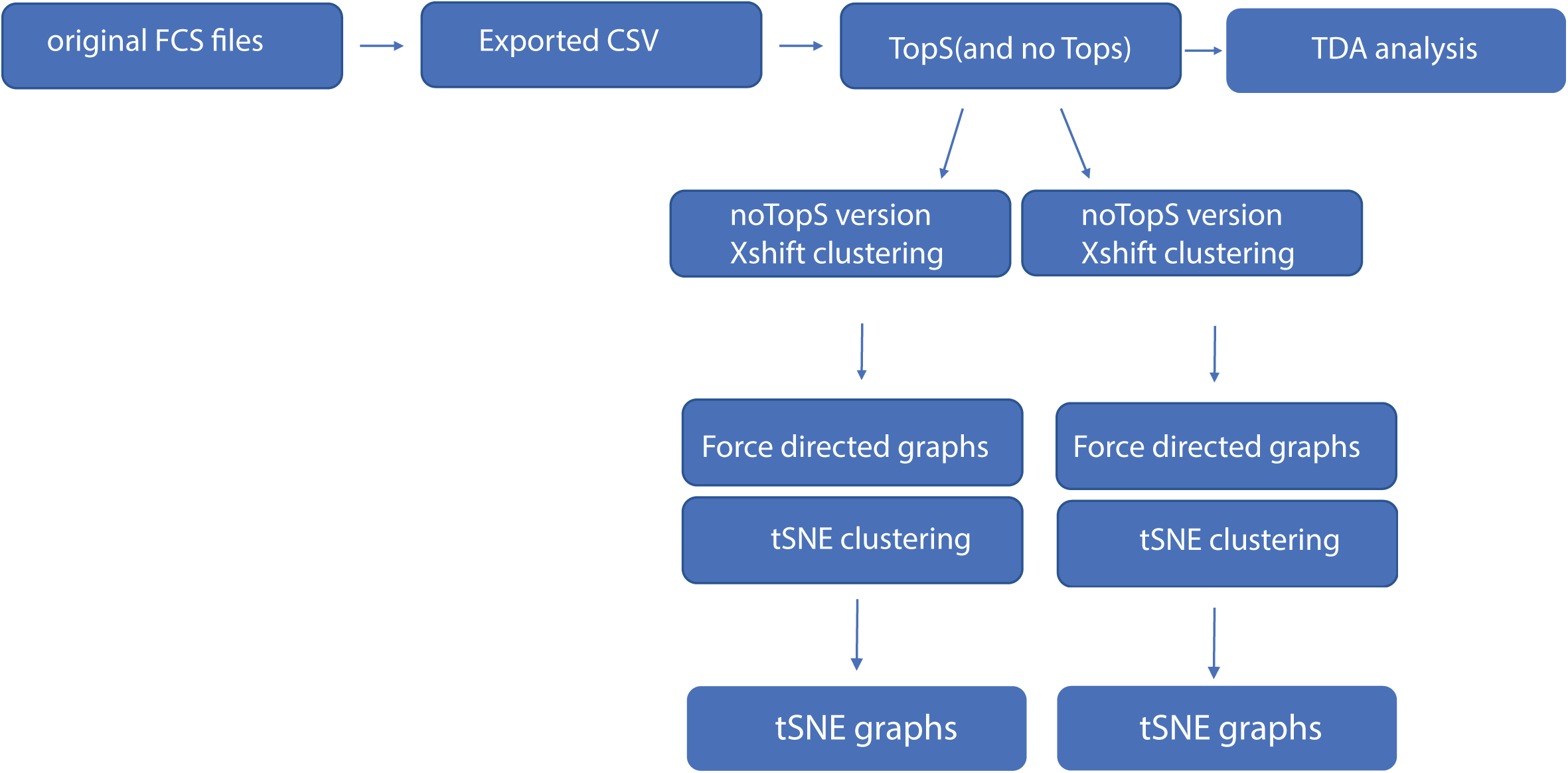
Overview of the computational flow. The computational approach started with the FSC files. The files were exported as CSV files. TopS shiny was used to produce the TopS values. TDA analysis was used for the original/transformed data and TopS values. The X-shift and t-SNE clustering were used for the original/transformed data and TopS values.

## Results and Discussion

### Clustering of human hematopoietic stem cell dataset

One of the major challenges for the study of hematopoietic stem cells (HSCs) is their identification and isolation from larger pools of cells ^9, 10^. Thus, developing biological and computational techniques for the identification of HSCs is of great importance. Here we selected a publicly available data from experiments in immunology using flow cytometry to demonstrate the use of TopS in the analysis of flow cytometry data. The *Nilsson rare* data contains rare population from bone marrow cells from healthy donor with 44,140 number of cells, 13 cell-surface protein markers and 358 (0.8%) manually gated cells (Supplementary Table 1) ^9, 10^. Early studies showed that no single cell-surface protein marker could specifically define the HSC and there is need of additional markers to purify HSC to homogeneity ^9, 10^. The *Nilsson rare* data consists of 13 different markers (i.e. CD10, CD110, CD11b, CD123, CD19, CD3, CD34, CD38, CD4, CD45, CD45RA, CD49fpur, CD90bio) that led to the identification of 9 different cell populations such as myeloid cells; B-lymphoid cells; CD4-T-cells; CD4+ T-cells; common lymphoid progenitors (CLPs); megakaryocyte/erythrocyte progenitors (MEPs); granulocyte/macrophage progenitors (GMPs); multipotent progenitor (MPPs); and hematopoietic stem cells (HSCs) ^9, 10^. The original data was pre-processed as described in Weber *et al*. ^10^ by using an arc-sinh transformation with a standard factor of 150 (i.e. *arcsinh*(x/150)) (Supplementary Table 1). From here on we call this matrix original/transformed data. TopS was next used to generate topological values on this dataset (Supplementary Table 2).

To better understand the changes in the expression of these cell-surface protein markers in the original/transformed data, we first applied a Pearson correlation (see Methods). In Figure 2A we represented the correlations between the cell-surface protein markers using their expression in the 44,140 cells. Overall, the Pearson correlations show a high range of correlations, ranging from rather low to high correlation coefficients. The highest correlations were between the CD110 and CD19 with a correlation of 0.911 followed by the correlation between CD19 and CD34 with a correlation of 0.857 (Fig. 2A). This result indicates that CD19, CD34 and CD110 might form a small cluster. In contrast, the lowest correlations were observed between CD3 and CD38 with an anticorrelation of -0.44147 followed by the correlation between CD10 and CD11b markers with an anticorrelation of -0.433 (Fig. 2A). These results suggest a substantial difference between the cell-surface protein markers profiles.

**Figure 2.**
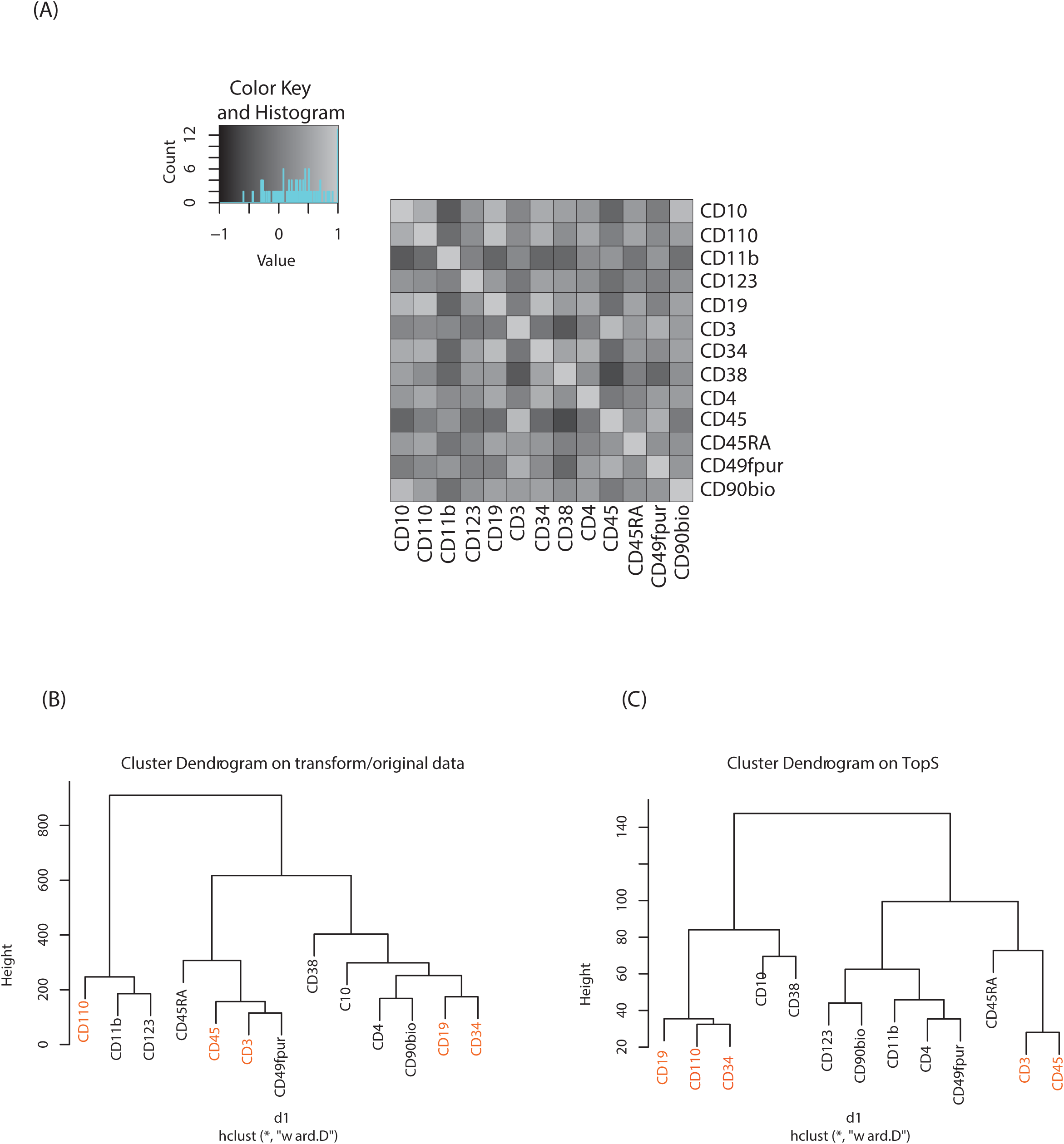
Pearson correlation and hierarchical clusters. (A) Pearson correlation was computed to show the similarity between cell-surface protein markers. For the original/transformed data (A) one hundred sixty-nine correlations are displayed in the figure. Hierarchical clustering was performed on the two matrices (i.e. original/transformed and Tops) with Euclidean distance and Ward as the method. In (B) and (C) the hierarchical clustering was performed on a 44,140 × 13 matrix using original/transformed data (B) and TopS values (C). The markers with the highest correlations in (A) were colored in red in (B) and (C). Note that in the shiny application the default parameters were set to Euclidian distance for the metric and Ward as the algorithm.

Hierarchical clustering was also performed on both the original/transformed data and the topological scores using the TopS shinny app to further illustrate the classification of the samples according to similarities of the cell-surface protein markers profiles (Fig. 2B). Interestingly, the markers pairs with the highest correlations are separated from each other when the original/transformed data is used (Fig. 2B). On the other hand, when using TopS the markers with the highest correlations in the matrix (i.e. CD19, CD110 and CD34) were under the same tree (Fig. 2C) in agreement with the Pearson correlations reported above. In addition, all the markers with the lowest correlations were positioned in both clusters away from each other (Fig. 2C). This figure illustrates the value of additional normalization methods like TopS to better elucidate the structure of the data and better cluster the samples. Furthermore, Figure 2 also suggested that various distance metrics must be explored when the transformed/original data is used.

### Topological scoring of dataset with machine learning approaches

Next, we utilized multiple machine learning approaches with or without TopS to evaluate the ability of TopS to discern rare cells in the *Nilsson rare* human hematopoietic stem cell dataset. To begin, we investigated the utility of topological data analysis (TDA) ^11-14^. TDA has be recently used for different omics data including cytometry data ^21^. TDA is a method that allows for the study of high dimensionality data sets by extracting shapes or patterns from the underlying data such that the researcher can gain new insights into patterns and relationships within the data ^11-14^. TDA also allows identification of clusters of rare events with a unique signature in a much larger data set ^11-14^. Because if its robustness to noise and coordinate insensitive nature and such has proved effective in identifying meaningful groupings or patterns of samples and data points from a diverse set of biological data types including microarray transcriptome data and protein-protein interaction networks ^22, 23^.

Here, the input data for TDA was represented in a matrix, with each column corresponding to each cell-surface protein marker and each row corresponding to a cell. The values were transformed values or topological scores for each cell-surface protein marker in different cell types. A network of nodes with edges between them was then created using the TDA approach based Ayasdi platform. Nodes in the network represent clusters of multiple cells, which is an important feature of the TDA network. This is in contrast to other networks where nodes consist of a single cell. Nodes in Figure 3 are colored based on the rows per node and on the label that corresponds to the gated cells (0 for major/multiple cells or 1 for HSC cells). Our aim was to provide a global overview of this complex dataset with the focus on the detection of rare events using TDA and additionally show the benefit of using TopS with TDA for the analysis of flow cytometry data. In Figure 3, we show the TDA analysis using (A) the topological score and (B) the original/transformed data in which the nodes are colored by the rows per node. In Figure 3A we observed that the cells are well separated in different groups based on the expression profiles. Importantly, Figure 3A also revealed group of cells in which the expressions of specific markers were enriched when compared with the rest of the markers, which is one of the unique features of the TopS. For example, we observed that the rare events were separated in two groups by TDA and the CD90bio and CD49fpur markers are enriched in these cells when compared with the other markers, and this agrees with the known association of CD90 and CD49f with human HSCs ^24^.

**Figure 3.**
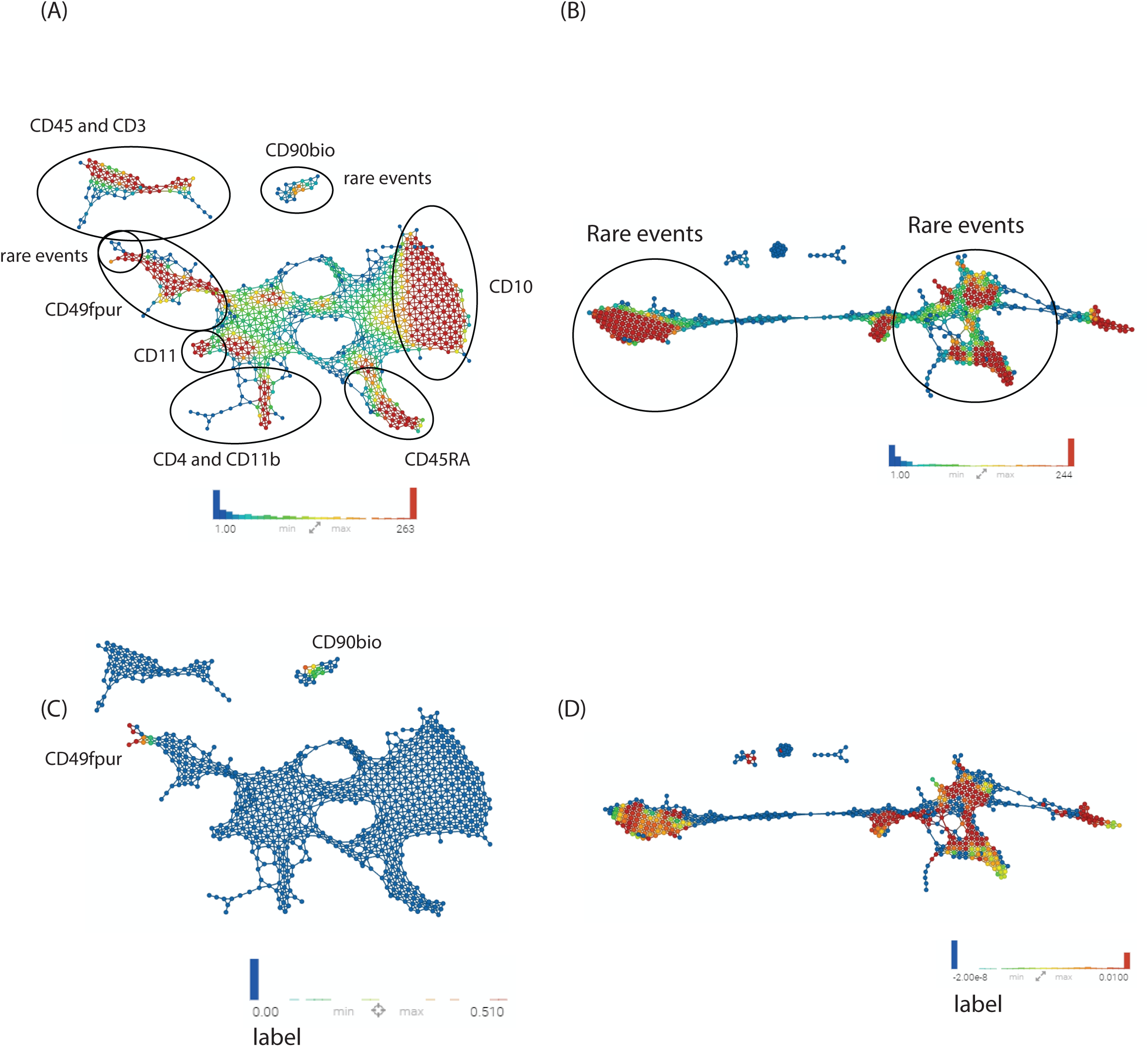
TDA data analysis. A TDA network was constructed for the original/transformed data and TopS scores. Correlation was used as a distance metric with 2 filter functions: Neighborhood lens 1 and Neighborhood lens2. Resolution 51 and gain 2 were used for 3A-3D. Node size is proportional with the number of cells in the node. Markers are illustrated in the figure. Cells are colored based on the rows per node for 3A and 3B. Color bar: red: high values, blue: low values. Cells were colored based on the label for the 3C and 3D. Cells were separated in different clusters based on their patterns in the 13 markers. The rare events were separated in two different clusters when using TopS values (A) and they were in multiple clusters when original/transformed data was used (B). The CD90, CD45 and CD49fpur markers were enriched in these rare events as shown in (A). The gated cells were colored in red, yellow or green (i.e. depending on the average per node) while the other cells were colored in blue. In 3C, we illustrated the rare events in the two clusters similar as in (A) while in (D) the rare events are spread throughout the network suggesting a poor separation.

TDA and TopS also detected other groups of cells where other markers were enriched. For example, of the right side of the Figure 3A, we can observe that the CD10 marker was highly expressed in the group of cells colored by red. TDA also shows a substantial amount of cross-talks between different markers. In contrast, in Figure 3B, when the original data/transformed data was used, TDA didn’t separate the data very well using the same parameters as in Figure 3A, and the majority of the rare events were spread through the entire network. To better highlight the location of the rare events in the two networks we colored the nodes by the label that corresponds to the gated cells and we observe a more focused localization of these cells when using TopS with TDA (Figures 3C and 3D).

We next investigated the use of X-shift clustering ^15, 16, 25, 26^ on the *Nilsson rare* flow cytometry data. X-shift (VorteX) is a standalone application with graphical interface that uses the weighted k-means density estimation ^15, 16, 25, 26^. Validation of the k value by elbow point gives an optimal k = 62 for 38 clusters in the case of TopS and k=62 for 30 clusters for the use of the original/transformed data (Supplementary Table 3). The results of TDA analysis using TopS data is in agreement with the results from the X-shift where the rare events were separated in two clusters. Similarly, X-shift produced two clusters for the rare events when the original/transformed data was used, however the overall numbers of clusters was smaller than the number of clusters obtained for TopS (Figure 4A and B, colored in blue). It is desirable to have more clusters than few in order to avoid smaller populations merging in larger clusters ^27^. Supplementary Figure 1A and 1B show that TopS provides wider range of numbers than in the original/transformed data, thus the over representative values in the matrix can be identified and therefore the markers that bring the most contribution in the detection of the rare events can be easily detected.

**Figure 4.**
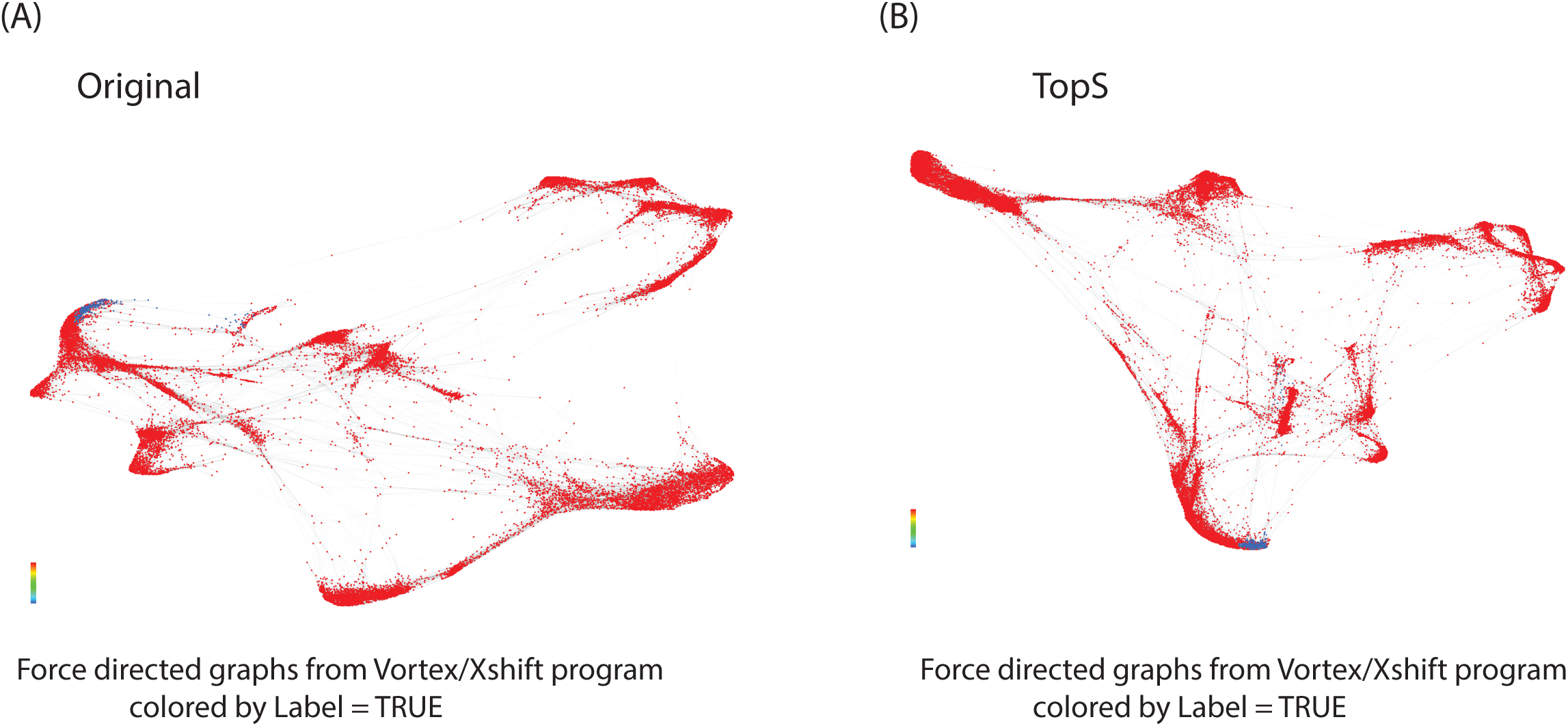
X-shift data analysis. X-shift(vortex) was used for the analysis of the original/transformed values and TopS values. Networks are colored by the label in Figure 4A and 4B. The gated cells are represented by the color blue.

Lastly, wet performed a t-SNE ^18-20^ analysis on the original/transformed and TopS data sets followed by a k-means clustering approach on the two vectors generated from the t-SNE (Supplementary Table 4). The number of clusters used for the k-means, were obtained from the X-shift as optimal numbers. Using k=38 for the TopS and k=30 in the case of the original/transformed data, the t-SNE produced similar results as the x-shift and TDA. Using TopS, the rare events were separated in two clusters (Figure 5A and 5B). Like X-shift, t-SNE recovered two clusters for the rare events when original/transformed data is used (Figure 5C and 5D). The smallest cluster identified cells in which the C90bio marker was remarkably expressed when compared with the other markers, while the largest cluster identified cells in which the C49fpur and CD45 were highly enriched (Supplementary Table 4). To visualize the difference between these two clusters we decided to represent the clusters as heat maps (Figure 6). The first cluster showed cells with high enrichment of several markers (Figure 6A) while the second cluster was an exception where cell populations has CD90bio with the highest enrichment (Figure 6B). These results also show that CD90bio, CD45 and CD49fpur are likely the most important markers among the 13 markers in the recovery of the HSC cells from this dataset ^9, 10^.

**Figure 5.**
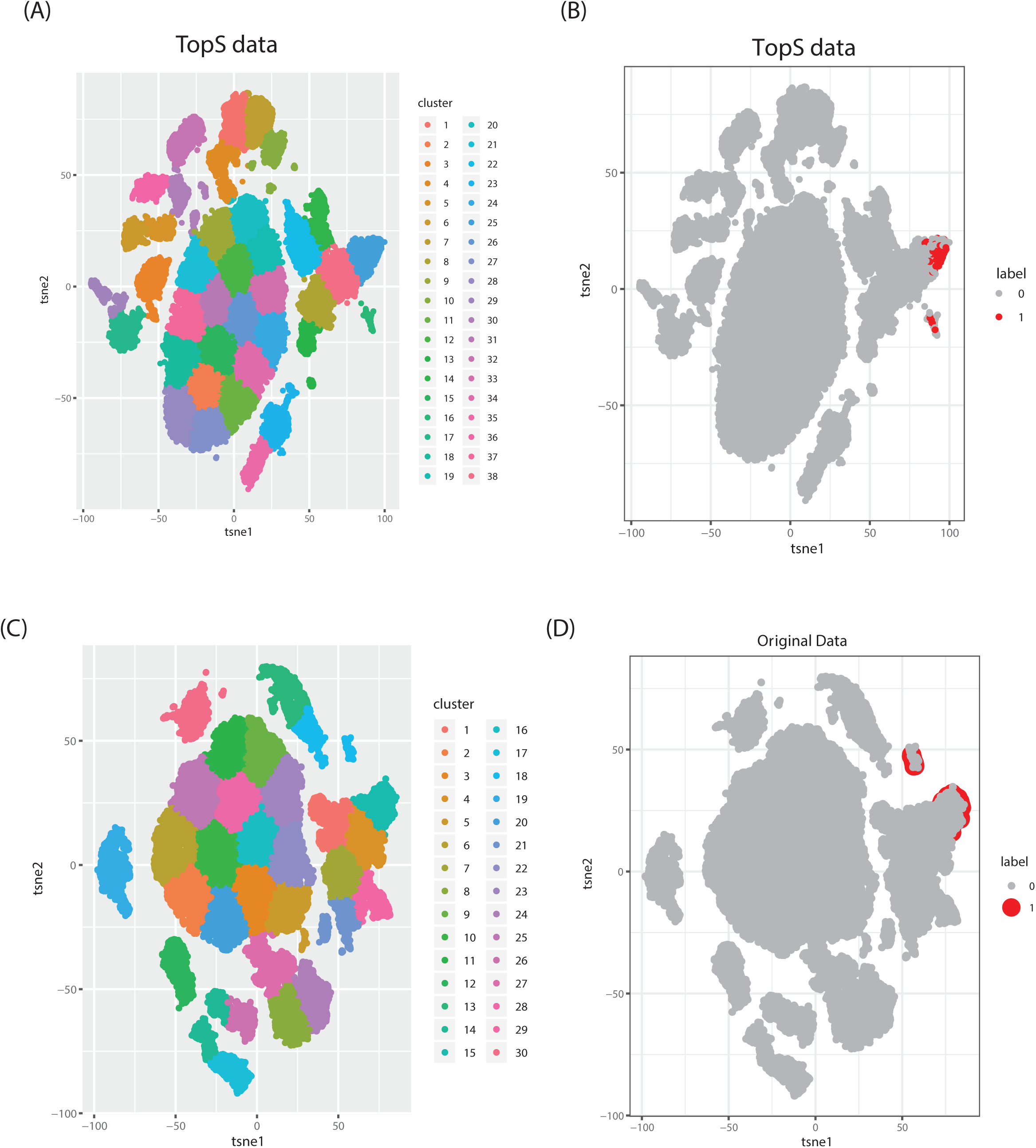
t-SNE data analysis. t-SNE analysis was implemented for the analysis of the original/transformed values and the TopS values. The networks were colored by the cluster numbers in Figure 5A and 5C. There are 38 clusters generated by t-SNE in (A) and 30 cluster in (C). In the case of Figure 5B and 5D, the networks are colored by the label. The gated cells are illustrated with the color red.

**Figure 6.**
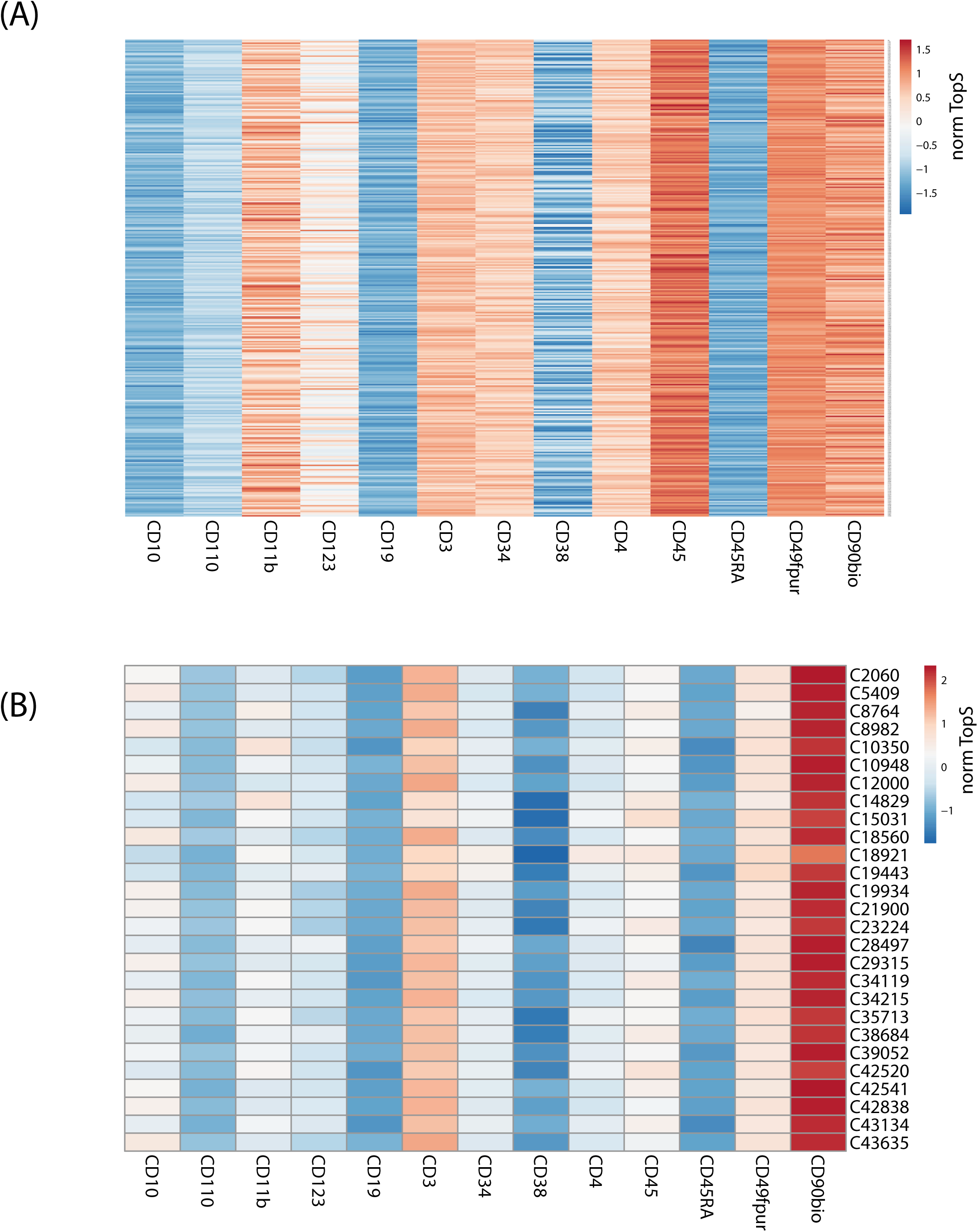
Heat maps. In Figures 6A and 6B we are illustrating only the gated cells which were separated in two different clusters by the t-SNE when TopS values were used. These figures show that some rare events had different patterns, i.e. cluster 1 shows different patterns than cluster. Consequently these have been separated in two clusters rather than grouped in a single cluster. In (A), we represented the gated cells in the first cluster where several protein markers were enriched in the HSC cells. High TopS values were observed in the C45, CD49fpur and CD90 in (A) as compared with other markers. In (B), we represented the gated cells in the second cluster (as determined by t-SNE) where the C90bio was highly express in these cells when compare with the other markers. The colors correspond to the TopS values, with blue representing low TopS values and red representing high TopS values. Heat maps were created using ClustVis software ^28^.

## Conclusions

High-dimensional flow cytometry is an important technique of choice to define and identify different population cells and detect expression levels of thousands of proteins markers ^9, 10^. We recently developed a topological score (i.e. TopS) that generates large range of values which subsequently can be used to identify overwhelming pairs, like those between an affinity purified protein and an associated protein, in a quantitative matrix ^5, 6^. Thus, subnetworks with highly scored pairs can be selected and visualized with TopS ^5, 6^. Previously, TopS was used on smaller matrices with thousands of rows for the analysis of networks, hence our goal was to extend its usage for the larger data sets like flow cytometry data. Here we tested the TopS shiny app for the analysis of a flow cytometry dataset described in Weber *et al*. ^10^ and we show the results for the *Nilsson rare* data ^9, 10^.

We first demonstrated that TopS values can be used with different clustering approaches for the analysis of the flow cytometry data. Using *Nilsson rare* data ^9, 10^, we applied three clustering methods with different approaches with the special focus on the identification of the rare events. Given the difficulties of identifying small clusters in a large dataset, TopS in combination with these methods identified the smallest population of rare events in a separate cluster (Figure 6B and Supplementary Figure 2). We demonstrated that rare populations have different patterns as they are pulled by different markers. As a result, they were separated in different clusters and not in a single cluster as one would expect. Using TopS we could identify a group of cells (Supplementary Figure 2 and Supplementary Table 4) in which the markers are having the highest expression. This data show that markers involved in T-cell and stem cells like CD11b, CD123, CD3 and CD90bio have the highest expressions in cells in this dataset.

However, when focusing only on the HSCs cells, we could show that TopS values revealed that the CD90bio, CD45, and CD49fpur are the most useful markers in the recovery of these cells and that a biological basis for the separation of HSCs into two clusters likely exists. These results could be beneficial for designing further experiments for the HSCs isolation. TopS in combination with machine learning can be effective in marker reduction (i.e from 13 markers to three/four markers) in the analysis of the bone marrow cells. Future work should focus on exploration of normalization methods and clustering approaches for a better representation of flow cytometry data. In conclusion, TopS ^5, 6^ could be an effective approach for processing flow cytometry data prior to further computational analysis with approaches like TDA ^11-14^, X-shift ^15-17^, and t-Distributed Stochastic Neighbor Embedding (t-SNE) ^18-20^.

## Experimental

### Data set

To evaluate the TopS method, we selected for our analysis an available data set from experiments in immunology using multicolor flow cytometry ^10, 25^. We have used a publicly available high-dimensional data set where cell population identities are known from the expert manual gating. The data was downloaded from the FlowRepository at https://flowrepository.org/id/FR-FCM-ZZPH. The data was manually gated cell population labels as the reference populations. The *Nilsson rare* data contain rare population from bone marrow cells from healthy donor with 44,140 number of cells, 13 cell-surface markers and 358(0.8%) manually gated cells ^10, 25^. Data was transformed using arcsinh and TopS method as described in *Sardiu et al*. ^6^. Pre-processing of the original data included the application of an arc-sinh transformation with a standard factor of 150 (i.e. *arcsinh*(x/150)).

### Overview of the TopS method

The TopS shiny app together with detailed documentation is freely available on github at https://github.com/WashburnLab/Topological-score-TopS-. Here, the raw data was assembled into a matrix in which the columns represent the individual cell-surface markers and rows represented pull-down cells. The elements of the matrix were represented by the transformed expression of cell-specific protein markers. The data was next normalized on the columns, rows and total sum of the numerical values in the matrix.

We used a simple model to calculate a score for each link between every cell and every marker in the matrix as follows:

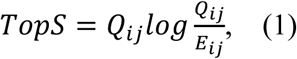

where Qij is the observed expression in row i and column j; and

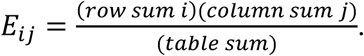

### Pearson correlation output

Pearson correlation was used here to illustrate the similarity between the 13 cell-protein markers (Figure 2A). Pearson correlation on the original/transformed data was calculated using R package cor(). The heatmap was used to illustrate the correlation between different samples using expression profiles. The heatmap shows a data matrix where coloring gives an overview of the numeric differences between cell-surface protein markers.

### Clustering with topological data analysis

The input data for TDA were represented in a matrix, with each column corresponding to each cell-surface protein marker and each row corresponding to a cell where the values are expression profiles. A network of nodes with edges between them was then created using the TDA approach based on the Ayasdi platform ^11-14^. Nodes in the network represent clusters of cells. Nodes in the figures were colored based on the metric Neighborhood lens 1 and Neighborhood lens2. Two types of parameters were needed to generate a topological analysis: The first is a measurement of similarity, called metric, which measures the distance between two points in space (i.e. between rows in the data). The second are lenses, which are real valued functions on the data points. Lenses could come from statistics (mean, max, min), from geometry (centrality, curvature) and machine learning (PCA/SVD, Autoencoders, Isomap). In the next step the data was partitioned. Lenses were used to create overlapping bins in the data set, where the bins are preimages under the lens of an interval. Overlapping families of intervals were used to create overlapping bins in the data. Metrics were used with <u>lenses</u> to construct the network output. There were two parameters used in defining the bins. One is *resolution*, which determines the number of bins. Higher resolution means more bins. The second is *gain*, which determines the degree of overlap of the intervals. Once the bins were constructed, we performed a clustering step on each bin, using single linkage clustering with a fixed heuristic for the choice of the scale parameter. This gives a family of clusters within the data, which may overlap, and we then constructed a network with one node for each such cluster, and we connected two nodes if the corresponding clusters contain a data point in common.

### Clustering with X-shift via Vortex

X-shift was running using graphical tool for cluster analysis of multiparametric datasets ^15, 16, 26^. The following parameters were used to run the X-shift application: transformation: none; noise threshold: yes, 1.0; feature rescaling: std; normalization: none; minimal Euclidean length: no; distance measure: angular distance; density estimate: N nearest neighbors; K from 150 to 5, steps 30; N: determine automatically; elbow point for automatic number of clusters was determined.

### Data analysis with t_SNE

To spatially map the cells in the dataset we first applied a t-distributed stochastic neighbor embedding(t-SNE), a nonlinear visualization of the data ^18-20^. We then applied k-means clustering to this transformed matrix using the Hartigan-Wong algorithm and a maximum number of iterations set at 50000. We used k=30 for the original/transformed data and k=38 for the TopS values to partition our data. The number of clusters were generated from the X-shift tool using elbow point. All computations were run using R environment using k-means function for the partition and daisy function to compute all the pairwise dissimilarities (Euclidean distances) between observations in the dataset for the silhouette.

## Acknowledgements

Research reported in this publication was supported by the Stowers Institute for Medical Research and the National Institute of General Medical Sciences of the National Institutes of Health under Award Number RO1GM112639 to MPW.

## Declaration of Interests

The authors declare no competing interests.

## Figure legends

**Supplementary Figure 1.**
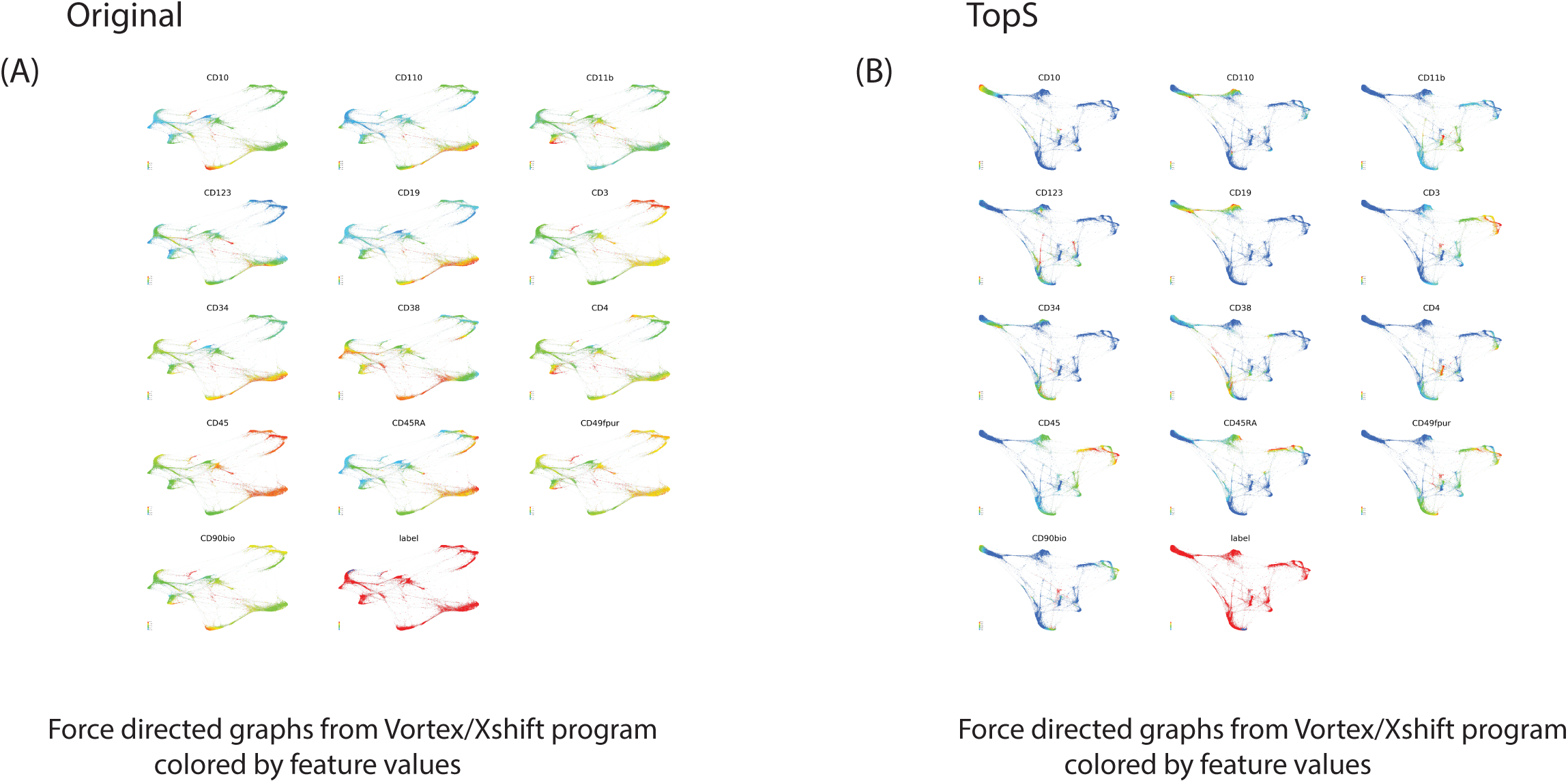
X-shift data analysis. X-shift(vortex) was used for the analysis of the original/transform values and TopS values. In (A) and (B) the networks are colored by the feature values. TopS provides wider range of numbers than in the than in the original/transformed data as shown by the color range. For example, TopS provides colors ranging from blue to red, while original/transformed values correspond to the colors ranging from green to red.

**Supplementary Figure 2.**
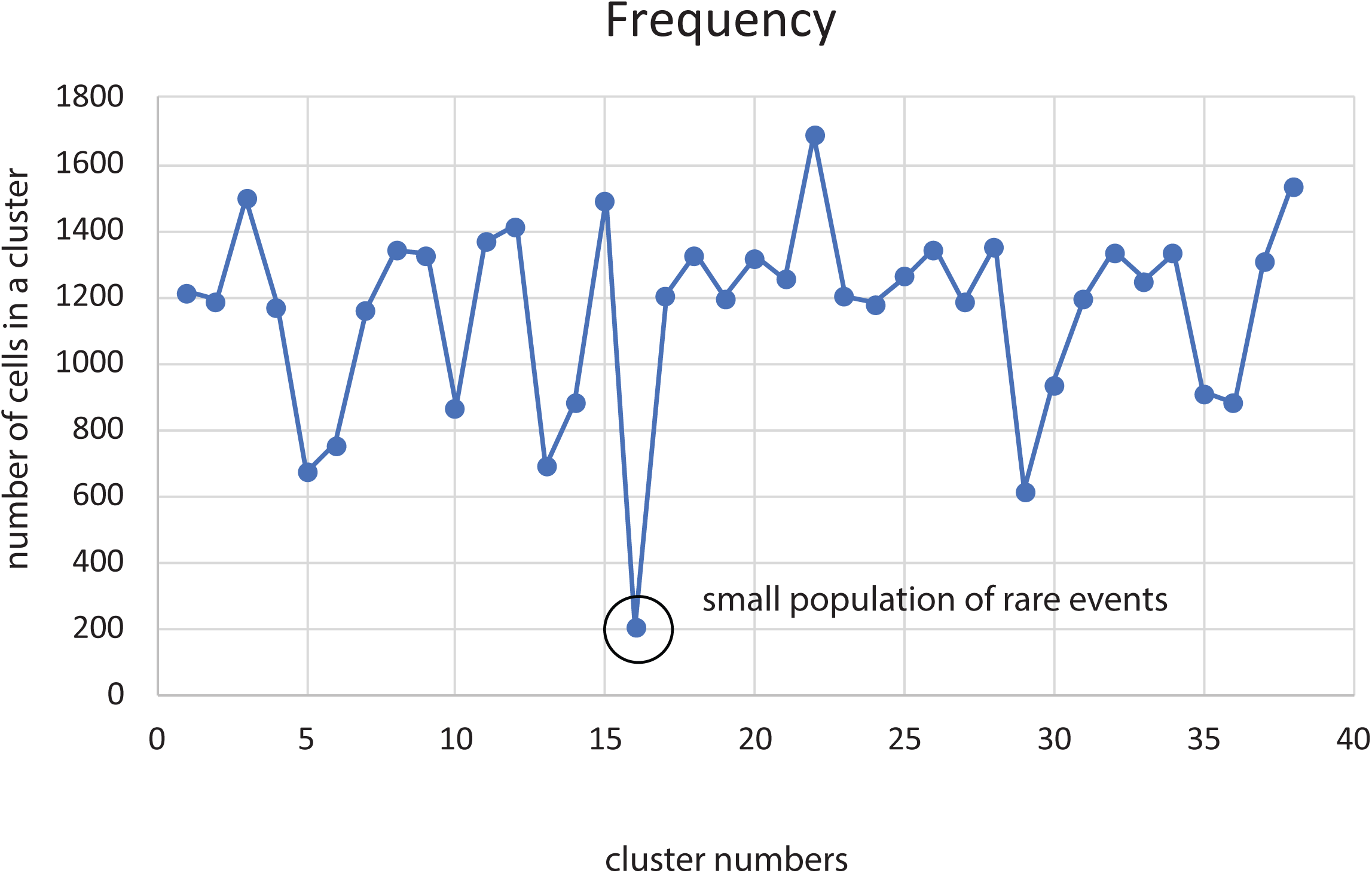
Clusters distribution. In this figure we illustrated the number of clusters detected by the t-SNE using TopS values and the number of cells in each cluster. t-SNE with TopS detected the smallest clusters consisting of gated cells with the marker C90bio enriched in these cells.

## References

1. C. Wu, F. Zhou, J. Ren, X. Li, Y. Jiang and S. Ma, High-throughput, 2019, 8.

2. Y. Li, F. X. Wu and A. Ngom, Briefings in bioinformatics, 2018, 19, 325–340.

3. Y. Hasin, M. Seldin and A. Lusis, Genome biology, 2017, 18, 83.

4. M. Bersanelli, E. Mosca, D. Remondini, E. Giampieri, C. Sala, G. Castellani and L. Milanesi, BMC bioinformatics, 2016, 17 Suppl 2, 15.

5. M. E. Sardiu, L. Florens and M. P. Washburn, Methods, 2019, DOI: 10.1016/j.ymeth.2019.08.010.

6. M. E. Sardiu, J. M. Gilmore, B. D. Groppe, A. Dutta, L. Florens and M. P. Washburn, Nature communications, 2019, 10, 1118.

7. M. J. Levy, D. C. Montgomery, M. E. Sardiu, J. L. Montano, S. E. Bergholtz, K. D. Nance, A. L. Thorpe, S. D. Fox, Q. Lin, T. Andresson, L. Florens, M. P. Washburn and J. L. Meier, Cell chemical biology, 2019, DOI: 10.1016/j.chembiol.2019.11.011.

8. G. Dayebgadoh, M. E. Sardiu, L. Florens and M. P. Washburn, Journal of proteome research, 2019, 18, 3479–3491.

9. A. Rundberg Nilsson, D. Bryder and C. J. Pronk, Cytometry. Part A : the journal of the International Society for Analytical Cytology, 2013, 83, 721–727.

10. L. M. Weber and M. D. Robinson, Cytometry. Part A : the journal of the International Society for Analytical Cytology, 2016, 89, 1084–1096.

11. P. G. Camara, Current opinion in systems biology, 2017, 1, 95–101.

12. P. G. Camara, D. I. Rosenbloom, K. J. Emmett, A. J. Levine and R. Rabadan, Cell systems, 2016, 3, 83–94.

13. L. Li, W. Y. Cheng, B. S. Glicksberg, O. Gottesman, R. Tamler, R. Chen, E. P. Bottinger and J. T. Dudley, Science translational medicine, 2015, 7, 311ra174.

14. P. Y. Lum, G. Singh, A. Lehman, T. Ishkanov, M. Vejdemo-Johansson, M. Alagappan, J. Carlsson and G. Carlsson, Scientific reports, 2013, 3, 1236.

15. M. Gossez, T. Rimmele, T. Andrieu, S. Debord, F. Bayle, C. Malcus, F. Poitevin-Later, G. Monneret and F. Venet, Scientific reports, 2018, 8, 17296.

16. N. Samusik, Z. Good, M. H. Spitzer, K. L. Davis and G. P. Nolan, Nature methods, 2016, 13, 493–496.

17. V. van Unen, T. Hollt, N. Pezzotti, N. Li, M. J. T. Reinders, E. Eisemann, F. Koning, A. Vilanova and B. P. F. Lelieveldt, Nature communications, 2017, 8, 1740.

18. N. V. Acuff and J. Linden, Journal of immunology, 2017, 198, 4539-4546.

19. A. Platzer, PloS one, 2013, 8, e56883.

20. S. Toghi Eshghi, A. Au-Yeung, C. Takahashi, C. R. Bolen, M. N. Nyachienga, S. P. Lear, C. Green, W. R. Mathews and W. E. O’Gorman, Frontiers in immunology, 2019, 10, 1194.

21. T. Lakshmikanth, A. Olin, Y. Chen, J. Mikes, E. Fredlund, M. Remberger, B. Omazic and P. Brodin, Cell reports, 2017, 20, 2238–2250.

22. M. E. Sardiu, J. M. Gilmore, B. Groppe, L. Florens and M. P. Washburn, Scientific reports, 2017, 7, 43845.

23. M. E. Sardiu, J. M. Gilmore, B. D. Groppe, D. Herman, S. R. Ramisetty, Y. Cai, J. Jin, R. C. Conaway, J. W. Conaway, L. Florens and M. P. Washburn, EMBO reports, 2015, 16, 116–126.

24. L. Lange, D. Hoffmann, A. Schwarzer, T. C. Ha, F. Philipp, D. Lenz, M. Morgan and A. Schambach, Stem cell reports, 2020, 14, 122–137.

25. J. Nilsson, I. Granrot, J. Mattsson, B. Omazic, M. Uhlin and S. Thunberg, Vox sanguinis, 2017, 112, 459–468.

26. A. K. Kimball, L. M. Oko, B. L. Bullock, R. A. Nemenoff, L. F. van Dyk and E. T. Clambey, Journal of immunology, 2018, 200, 3–22.

27. G. K. Chen, E. C. Chi, J. M. Ranola and K. Lange, PLoS computational biology, 2015, 11, e1004228.

28. T. Metsalu and J. Vilo, Nucleic acids research, 2015, 43, W566–570.

